# Optogenetic suppression of corticogeniculate feedback in anesthetized ferrets is overridden by visual stimulation

**DOI:** 10.1101/2021.06.28.450254

**Authors:** S. Zhu, J.M. Hasse, F. Briggs

**Affiliations:** Neuroscience Graduate Program, University of Rochester Medical Center, Rochester NY 14642 USA; Center for Neural Science, New York University, New York, NY 10003 USA; Department of Neuroscience, University of Rochester School of Medicine, Rochester NY 14642 USA; Department of Brain and Cognitive Sciences, University of Rochester, Rochester NY 14627 USA; Ernest J. Del Monte Institute for Neuroscience, University of Rochester School of Medicine, Rochester NY 14642 USA; Center for Visual Science, University of Rochester, Rochester NY 14627 USA

## Abstract

The feedforward projection from the retina shapes the spatial receptive field properties of neurons in the dorsal lateral geniculate nucleus of the thalamus (LGN). Corticogeniculate feedback from the visual cortex appears to exert a more subtle, modulatory influence on LGN responses. Studies involving manipulations of corticogeniculate feedback have yielded inconsistent findings, but the reasons for these inconsistencies are not known. To examine the functional contributions of corticogeniculate feedback, and to resolve past inconsistencies, we examined the effects of selective optogenetic suppression of corticogeniculate neurons in anesthetized ferrets. In particular, we examined the responses of LGN and V1 neurons during optogenetic suppression of corticogeniculate feedback in the presence and absence of visual stimulation and across conditions in which the frequency of LED illumination varied. Optogenetic suppression of corticogeniculate feedback decreased activity among LGN neurons in the absence of visual stimulation, dispelling the notion that anesthesia causes a floor effect. In contrast, suppressing corticogeniculate feedback did not affect the visual responses of LGN neurons, suggesting that feedforward visual stimulus drive overrides weak corticogeniculate influence. Optogenetic effects on LGN and V1 neuronal responses depended on the frequency of LED illumination, with higher frequency illumination inducing slow oscillations in V1, dis-inhibiting V1 neurons locally, and producing more suppression among LGN neurons. These results demonstrate that corticogeniculate influence depends on stimulation parameters including visual stimulus conditions and frequency of inactivation. Furthermore, weak corticogeniculate influence is overridden by strong feedforward visual stimulus drive – this attribute is the most likely source of inconsistencies in past studies.

**Significance Statement:** Although corticogeniculate synapses onto thalamic neurons far outnumber those coming from the retina, the function of corticogeniculate feedback in vision has remained a stubborn puzzle. Prior studies of corticogeniculate feedback have yielded inconsistent findings, but the source for these inconsistencies is unknown. We utilized selective optogenetic suppression of corticogeniculate feedback to examine its effects on thalamic neuronal responses and to resolve sources of prior inconsistencies. We found that suppression of corticogeniculate feedback reduced thalamic responses, but only in the absence of visual input. This suggests that the major source of inconsistencies across prior studies is the fact that weak corticogeniculate influence is overcome by strong feedforward visual stimulus drive.

## Introduction

In the visual system, the functional significance of feedforward projections from the retina to the dorsal lateral geniculate nucleus of the thalamus (LGN) and from the LGN to the primary visual cortex (V1) are well established (Sherman & Guillery 2006). However, corticogeniculate feedback to the LGN, which mainly comes from V1 (see for review, Briggs 2020), is less well understood. Curiously, corticogeniculate feedback contributes more than 30% of the total synaptic inputs onto LGN neurons, while feedforward inputs from the retina contribute less than 10% (Erisir et al 1997, Van Horn et al 2000). Its numerically robust anatomical input onto LGN neurons suggests that corticogeniculate feedback plays an integral role in visual processing.

Corticogeniculate feedback affects various aspects of LGN activity including response gain (Hasse & Briggs 2017, Przybyszewski et al 2000, Webb et al 2002), spatiotemporal resolution (Andolina et al 2013, Hasse & Briggs 2017, Webb et al 2002), inter-neuronal synchronization (Sillito et al 1994, Weliky & Katz 1999), contrast adaptation (Li et al 2011), gain variability (Murphy et al 2020a), and information coding (McClurkin et al 1994, Murphy et al 2020a). However, these effects are often weak and inconsistent across studies, leading to a lack of consensus on the functional role of corticogeniculate feedback in vision. For example, Przybyszewski et al (2000) showed a reduction of visual response gain in the majority of LGN neurons after inactivating V1 with cortical cooling in anesthetized macaque monkeys. Webb et al (2002) lesioned V1 in anesthetized marmosets and found reduced response gain only in magnocellular cells in the LGN. More diverse effects on LGN visual responses were observed by Wang et al (2018) following pharmacological activation of V1, by Denman and Contreras (2015) following optogenetic inactivation of corticogeniculate neurons, and by Geisert et al (1981) following V1 cooling: all groups observed increases, decreases, and no change in response gain among LGN neurons as a result of their experimental manipulations.

Inconsistent findings across studies could arise from the complexity of corticogeniculate circuitry. It is well-established that the feedforward pathways from the retina to V1 are segregated into multiple parallel processing streams with different morphological and physiological properties: parvocellular, magnocellular and koniocellular streams in primates, and X, Y and W streams in carnivores (Kaplan 2014, Sherman & Guillery 2006). Corticogeniculate feedback is also organized into parallel streams that align with the feedforward streams (Briggs & Usrey 2005, Briggs & Usrey 2009). In addition to making monosynaptic excitatory connections onto LGN relay neurons, corticogeniculate neurons also disynaptically inhibit LGN relay neurons via neurons in the thalamic reticular nucleus (TRN) and local LGN inhibitory neurons (Sherman & Guillery 2006). Accordingly, corticogeniculate feedback could differentially regulate information transfer in each of the parallel streams and/or through differential regulation of excitatory and inhibitory inputs. On the other hand, methodological differences could also contribute to inconsistent findings across studies. First, most studies utilized procedures to irreversibly inactivate corticogeniculate feedback in anesthetized animals. Since corticothalamic activity is reduced by anesthesia (Steriade 2003), this may have generated a floor effect, which could not be estimated following an irreversible manipulation. Second, most studies utilized non-selective manipulations that affected the visual cortex broadly. Third, most previous studies did not consider the possibility that corticothalamic feedback influence may depend on the firing mode of corticothalamic neurons, which can result in net facilitation or suppression of thalamic activity (Crandall et al 2015, Kirchgessner et al 2020). Fourth, most studies investigated the function of corticogeniculate feedback while the feedforward visual pathways were activated by visual stimuli. Activation of strong feedforward inputs from the retina that shape the spatial receptive field properties of LGN neurons (Usrey et al 1999) may have obscured subtle modulatory effects of corticogeniculate feedback (but see Hasse & Briggs 2017). Overall, the extent to which particular factors such as anesthesia, frequency or selectivity of corticogeniculate manipulation, or overriding feedforward stimulus drive confounded examinations of corticogeniculate function remained unresolved.

Here, we examined whether selective and reversible optogenetic suppression of corticogeniculate feedback alters the visual response properties of LGN and V1 neurons and whether specific confounding experimental factors are responsible for inconsistencies regarding corticogeniculate function. We performed these studies in anesthetized ferrets, diurnal predators with good vision (Jackson & Hickey 1985). Suppressing corticogeniculate feedback reduced LGN responses, but only in the absence of visual stimulation. Furthermore, effects of optogenetic suppression of corticogeniculate feedback depended on the frequency of LED illumination. Overall, the effects of corticogeniculate suppression in the LGN were weak and easily overcome by feedforward visual stimulus drive.

## Methods and Materials

### Animals

Data for this study were obtained from three adult female ferrets (*Mustela putorius furo*). All animal procedures were approved by the Institutional Animal Care and Use Committee at the research institution and adhered to the guidelines for animal use issued by the US Department of Agriculture and NIH.

### Viral injections

About 5μl of SADΔG-ArchT-GFP rabies virus, obtained from Dr. Fumitaka Osakada working in the laboratory of Dr. Ed Callaway at the Salk Institute, was injected into the ferret LGN during a surgical procedure performed under full surgical anesthesia and employing aseptic techniques. Virus injection methods followed those described previously (Hasse & Briggs 2017). The location and depth of the LGN were determined neurophysiologically and following stereotaxic coordinates. The virus was loaded in a glass pipette and injected into the LGN at the designated location and depth using a Nanoject (Drummond Scientific Company). The virus infected corticogeniculate neurons through their axon terminals in the LGN, as has been documented previously for similar rabies virus manipulations (Briggs et al, 2016; Hasse & Briggs, 2017; Hasse et al, 2019; Murphy et al, 2020). This particular virus expressed the inhibitory optogenetic proton pump archaerhodopsin (ArchT) and GFP in infected neurons. In two of three ferrets, the virus successfully infected corticogeniculate neurons. These are referred to as Experimental animals throughout. In the third ferret, referred to as the Control animal, no virus infection was detected, which is equivalent to sham injection.

### Electrophysiology and optogenetics

Electrophysiological recordings were done 7-8 days post injection. A 24-contact linear multielectrode array (U-Probe from Plexon Inc., Dallas, TX) was inserted into V1 and an array of seven independently movable platinum-tungsten microelectrodes (Eckhorn Matrix from Thomas Recording GMBH, Giessen, Germany) was inserted into the LGN to record extracellular spikes simultaneously from both brain areas in anesthetized and paralyzed ferrets. Receptive fields of recorded neurons were first mapped using standard hand-mapping and software-assisted procedures. Drifting gratings varying in contrast (0-100%), temporal frequency (TF; 1-32Hz), spatial frequency (SF; 0.1-2.5 cycles/degree), orientation (0-324 degrees), or size (0.32-12.6 degrees) in steps of 10 were then presented in the receptive fields of recorded neurons. Flashing black spots (100 msec flash, 2Hz cycle) 12-30 degrees in diameter placed within recorded neuronal receptive fields were used to measure neuronal response latencies. White noise m-sequence stimuli (square grid 15-30 degrees across) were placed within recorded neuronal receptive fields to measure spatiotemporal receptive fields. Spontaneous activity was measured during periods when the monitor displayed mean grey. In half of all trials, corticogeniculate feedback was suppressed by shining a green LED (515 nm; ~0.5-2mW/mm^2^ maximum) on the surface of V1. To quantify optogenetic suppression of LGN and V1 activity in the absence of visual stimulation, the LED was flashed at 2Hz or 4Hz continuously (in one Experimental and the Control animal), or at 0.5-16.5Hz in steps of 10 (in one Experimental animal) with 2s periods of flashes at each frequency followed by 2s inter-flash intervals. LED flashes paired with visual stimuli were delivered in synch with and at the same temporal frequency as drifting gratings or flashing spots. LED flashes rose and decayed within 6 msec, so they followed the highest temporal frequency used (32 Hz). For LED flash frequencies above 8Hz, there was a roughly linear increase in LED power measured at the surface (from 0.1 to 0.5mW). The LED was continuously on during white noise m-sequence stimulation. All visual stimulus sequences were repeated 2 or 3 times, i.e. 2 or 3 repeats without LED stimulation and 2 or 3 repeats with LED stimulation. LED-only stimulation, in the absence of visual stimulation, was repeated twice. Spike sorting of single units was done manually using standard principal components analysis (PCA) based analyses within a commercial software package (Offline Sorter from Plexon Inc., Dallas, TX). V1 and LGN multiunits were obtained by manually setting a threshold for each channel to exclude noise, which was held consistent across recordings per channel, and selecting all threshold crossings as multiunit spikes.

### Histology

Viral infection of corticogeniculate neurons was verified for all animals using histological analyses. At the end of the electrophysiological recording experiment, animals were euthanized and perfused transcardially. The brain was extracted and cryo-sectioned. Coronal sections of the LGN and V1 were first stained for cytochrome oxidase activity to visualize layers. To permanently stain infected neurons, sections were labeled with a primary antibody against GFP (A-11122, Thermo Scientific, Waltham, MA), and with a biotinylated secondary antibody (goat anti-rabbit, Molecular Probes/Life Technologies, Grand Island, NY), before being reacted with DAB/peroxide. Numbers of virus-infected neurons per histological section containing area 17 were counted using a Neurolucida camera and software package (MicroBrightField, Williston, VT).

### Data inclusion

We analyzed 8 LGN and 4 V1 single units and 10 LGN and 17 V1 multiunits from Experimental animal I, and 16 LGN single units and 23 V1 multiunits from Experimental animal II. From the Control animal, 19 LGN single units were analyzed, however, one recording penetration with 5 single units was removed from further analyses because it was incomplete. Therefore, we analyzed a total of 24 LGN single units, 4 V1 single units, 10 LGN multiunits and 40 V1 multunits from two Experimental animals and 14 LGN single units from the Control animal. We only analyzed V1 multiunits recorded from electrode contacts that were approximately 0.8 – 2 mm below the brain surface in order to target deep layer V1 neurons, including putative virus-infected corticogeniculate neurons. All of the single units (from here onward referred to as ‘neurons’) analyzed satisfied our inclusion criteria of good spike isolation based on PCA clustering with interspike interval violations less than 1%. From the two Experimental animals, 23 of 24 LGN neurons and all the V1 neurons displayed good tuning (i.e. R square of curve fitting larger than 0.5) in both LED-ON and LED-OFF conditions in at least one visual stimulus test (e.g. contrast, SF, TF, or size). From the Control animal, 6 of 14 LGN neurons displayed good tuning. For analysis of LGN activity during optogenetic suppression of corticogeniculate neurons in the absence of visual stimulation (slope and ABC analyses described below), data were available for 21 LGN neurons from the two Expeirmental animals and from all LGN neurons from the Control animal. Two LGN neurons from Experimental animal II and 3 LGN neurons from the Control animal were excluded from the slope and ABC analyses due to too low (< 3 Hz) mean firing rates during the LED-ON period. For the ABC analysis, 1 LGN neuron from Experimental animal I, 3 LGN neurons from Experimental animal II, and 5 LGN neurons from the Control animal were excluded due to too low (< ½ of the mean firing rate during LED) or high (> 2 times of the mean firing rate during LED) baseline firing rates that made ABC values uninterpretable. Of the 19 LGN neurons from Experimental animals used for the slope analysis, 13 LGN neurons from Experimental animal I contributed two data points. Of the 11 LGN neurons from the Control animal used for the slope analysis, 5 LGN neurons contributed two data points. Of the 15 LGN neurons from Experimantal animals used for the ABC analysis, 6 LGN neurons from Experimental animal I contributed two data points. Of the 7 LGN neurons from Control animals used for the ABC analysis, 2 LGN neurons contributed two data points. Contributions of two data points per LGN neuron were possible because data were available for two different LED conditions: with 2Hz and 4Hz LED flashes (**Figures 2C** and **D**). Numbers of LGN neurons from the two Experimental animals with tuning for each visual stimulus test are indicated in **Table 1**. Although all 23 LGN neurons from Experimental animals with some tuning were included in **Table 1**, if a neuron had poor tuning for a given metric in the LED-OFF or LED-ON condition, tuning data for that metric and condition were excluded. Numbers of LGN neurons from the two Expeimental animal included in the PSTH slope and AUC analyses are indicated in **Table 2**. LGN neurons were excluded from the PSTH slope analysis for a given tuning metric and both LED conditions if the firing rate was low or if neurons were poorly tuned in either LED condition. LGN neurons were excluded from the AUC analysis for a given metric and both LED conditions if neurons were poorly tuned in either LED condition. From Experimental animals, responses to varying LED frequencies were available for 16 V1 multiunits, which were included in the power spectral density analysis described below.

**Table 1:**
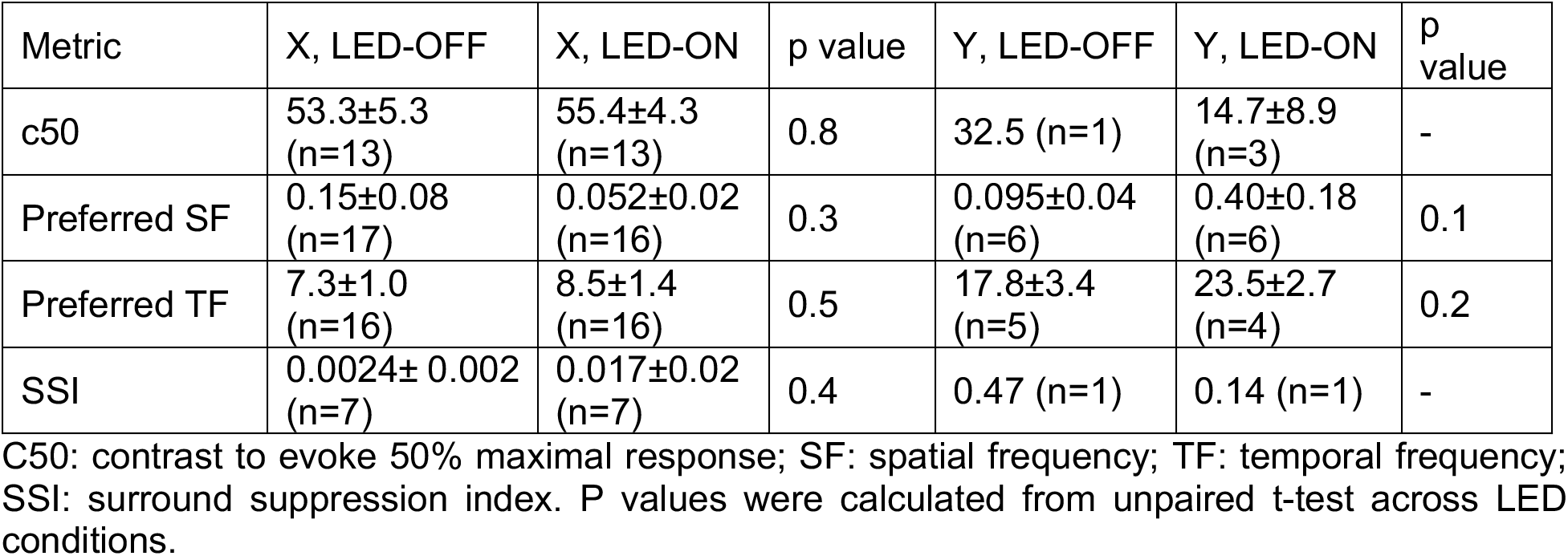
Effects of optogenetic suppression of corticogeniculate neurons on LGN tuning

**Table 2:**
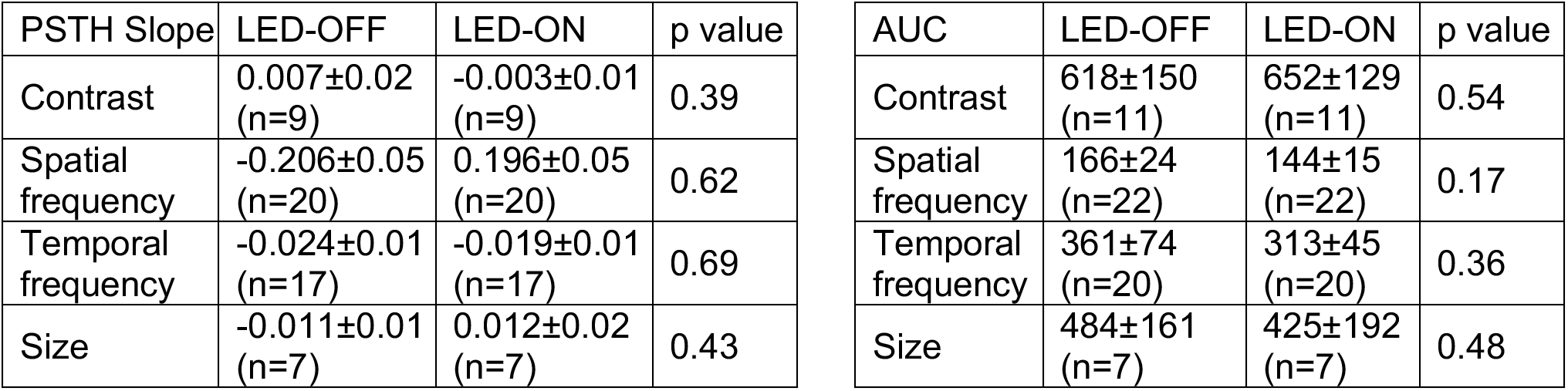
Effects of optogenetic suppression of corticogeniculate neurons on LGN visual PSTH slope and AUC. P values were calculated from paired t-test across LED conditions.

### Tuning measurements

To generate tuning curves, the firing rates of LGN and V1 neurons and V1 multiunits in response to gratings varying in contrast, TF, SF, orientation, or size were baseline-corrected by subtracting the averaged firing rate during the 2s inter-trial period of each trial between stimulus presentations. Tuning responses were averaged across the 2-3 repetitions per visual stimulus test and LED condition. Contrast responses were fitted with 1^st^ or 2^nd^ order power functions; SF, TF, and size responses were fitted with 1^st^, 2^nd^, or 3^rd^ order Gaussian functions. The contrast to evoke a half-maximal response (c50) was computed per LGN neuron, V1 neuron, and V1 multiunit from curve fits. Additional tuning metrics computed per LGN neuron, V1 neuron, and V1 multiunit included the preferred SF and TF corresponding to the global peak of the tuning curve fits and the surround suppression index (SSI) computed as the difference between the curve fit maximum and the plateau, corresponding to an easing of the slope of the size tuning curve fit past the peak, divided by their sum.

### Generating spike density functions (SDFs) and peri-stimulus time histograms (PSTHs)

For the analysis of neuronal responses to optogenetic suppression of corticogeniculate neurons in the absence of visual stimulation, SDFs with a square kernel were generated for each neuron by sliding a 250ms window across the recording period (total duration = 37s) in steps of 50ms, and calculating the firing rate (spike per second) within each window (Szucs 1998). The firing rate was then normalized by the mean firing rate during the LED-ON period per neuron. For all neurons, activity recorded between 250ms after the start of the recording and 2s after the LED onset was excluded from the SDF calculations due to an unexpected monitor screen flicker that occurred on a proportion of trials during this time window. For neuronal responses to visual stimulation, standard PSTHs of individual neuronal activity were generated using a bin size of 50ms, and were then normalized by the mean firing rate during visual stimulus presentation per neuron. Average SDFs and PSTHs were generated by averaging the normalized SDFs or PSTHs across all LGN neurons in the sample. For display purposes only, the average SDF in Figure 2 and the average PSTH in Figure 3 were smoothed.

### Quantification of the effect of optogenetic suppression of corticogeniculate feedback on LGN activity without and with visual stimulation

For all quantifications of LGN responses to LED modulation, a baseline firing rate was calculated from the first 250ms of the recording, during which time there was no LED illumination or visual stimulation. For LGN neuronal responses to LED modulation in the absence and presence of visual stimulation, a straight line was fitted to the first 15s of each neuronal SDF or PSTH, respectively, and the slope of that line was calculated. The more negative the slope, the more suppression of LGN activity occurred following LED onset. For LGN responses to LED modulation in the absence of visual stimulation, the area between the baseline firing rate extrapolation line and the neuronal SDF was computed over the full duration of LED presentation (denoted as the area between the curves or ABC). The more negative the ABC, the more suppression of LGN activity during LED presentation. For 8 LGN neurons and an additional 10 LGN multiunits, firing rates during LED illumination in the absence of visual stimulation were computed separately to responses to low (<2Hz) and high (>8Hz) frequency LED flashes. For LGN responses to LED modulation in the presence of visual stimulation, a separate quantification was performed. The area under the baseline-corrected tuning curve (AUC) was computed to quantify the magnitude of the visual response, which was then compared across trials with and without LED illumination. The more positive the AUC, the greater the visual response magnitude.

### Quantification of response latency and precision

We used reverse correlated m-sequence frames to generate spatiotemporal maps of the spike-triggered average (STA) of LGN responses to the white noise stimulus. The frame with the peak pixel brightness (strongest light/dark response for ON/OFF LGN neurons compared to background) was used to determine the extent of the spatial receptive field using a 2-dimensional Gaussian fit. The brightness of the peak pixel within the fit was then tracked across STA frames and the pixel brightness over time data were fitted with a 2^nd^ order Gaussian function to generate the pixel brightness curve. The latency was the time corresponding to the maximum/minimum of this curve for ON/OFF LGN neurons, respectively, relative to the time of the spike. The spike timing precision of LGN neurons was evaluated from their responses to the flashing spot stimulus. An event start time was manually defined for each LGN neuron to discount spontaneous activity before the neuronal response to the flashed stimulus. We then calculated the standard deviation of the time of the first spike after this event start time. We also measured the latency of LGN neuronal responses to the flashing spot stimulus by computing the time to the half-maximal firing rate in the flash-evoked PSTH.

### Power spectral density (PSD) analysis

The PSD analysis was used to examine oscillations in activity among 4 deep layer V1 neurons and 16 V1 multiunits from Experimental animal II in which the LED was flashed at both low and high frequencies. Separate SDFs computed during low-frequency (0.5-1.5Hz) or high-frequency (8.5-16.5Hz) LED flashes were averaged across the 2 repetitions per V1 unit. SDFs were fit with smoothing splines, which were then used to calculate the power between 0.15 and 0.6Hz using Welch’s method (‘pwelch’ function in MatLab) with a Hamming window of 12s, the duration of each SDF. Power spectra per V1 unit were normalized to the averaged power between 0.15 and 0.6Hz per V1 unit. Normalized power spectra were plotted for each V1 unit along with average power spectra across all V1 units separately for low and high LED flash frequency responses. The peak frequencies per V1 unit for low and high LED flash frequency responses were computed as the frequency corresponding to the maximum of each power spectrum.

### Software accessibility

Custom Matlab code was written to perform the data analyses outlined above. The authors will make this code available upon request.

## Results

There are a number of reasons why it has been difficult to pinpoint the function of corticogeniculate feedback in visual perception. Inconsistent findings across studies may be partially attributed to methodological differences, however some variation in experimental outcomes could be attributed to the fact that corticothalamic neurons are suppressed during slow cortical oscillations in anesthesia and sleep (Steriade 2003). Accordingly, experimental manipulations that suppress corticogeniculate feedback in anesthetized preparations may run into a floor effect that is difficult to assess with irreversible manipulation of feedback. Other confounding factors leading to inconsistent findings include the fact that corticogeniculate circuits in highly visual animals are heterogeneous (Briggs et al 2016, Briggs & Usrey 2005, Briggs & Usrey 2009, Hasse et al 2019) and corticogeniculate influence on LGN responses may depend on the firing mode of corticogeniculate neurons (Kirchgessner et al 2020). Alternatively, because corticogeniculate feedback merely modulates LGN responses (Guillery & Sherman 2002, Sherman & Guillery 1998), it is possible that visual stimulus drive from the retina overrides the subtle impact of experimental suppression of corticogeniculate feedback on LGN responses. To test these alternatives explicitly using selective and reversible manipulation of corticogeniculate neurons, we measured LGN neuronal responses during optogenetic suppression of corticogeniculate feedback in the presence and absence of visual stimulation in anesthetized ferrets.

To achieve selective and reversible optogenetic suppression of corticogeniculate neurons, a glycoprotein(G)-deleted rabies virus (SADΔG-ArchT-GFP) was injected into the LGN of three ferrets. In two Experimental ferrets, virus injection into the LGN was successful in driving the expression of optogenetic proteins in corticogeniculate neurons (**Figure 1**), while in one Control ferret, there was no viral infection of any V1 neurons and data from this animal were utilized for control comparisons. The virus infected corticogeniculate neurons through their axon terminals in the LGN and drove expression of the inhibitory opsin Archaerhodopsin (ArchT) and GFP (**Figure 1A** and **C**). All virus-infected and labeled neurons in V1 were located in layer 6, confirming that the virus specifically infected corticogeniculate neurons. **Figure 1B** and **D** shows virus injection zones in two Experimental animals with darkly stained viral particles and sparse labeling of LGN neurons. It is interesting to note that in this study, as well as in previous studies (Hasse & Briggs 2017, Murphy et al 2020b), few virus-infected LGN neurons were observed, suggesting low transduction efficiency for LGN interneurons. Although the total numbers of virus-infected corticogeniculate neurons were not high per Experimental animal, labeled neurons were present throughout area 17 of each Experimental animal (**Figure 1E**). Seven to eight days post virus injection, ferrets were anesthetized and paralyzed for electrophysiological recordings. LGN activity was recorded using an array of seven independently movable platinum-tungsten microelectrodes, and V1 activity was recorded using a 24-contact linear electrode. **Figure 1C** shows a lesion from an electrode, demonstrating that recording sites in V1 were close to virus-infected corticogeniculate neurons.

**Figure 1:**
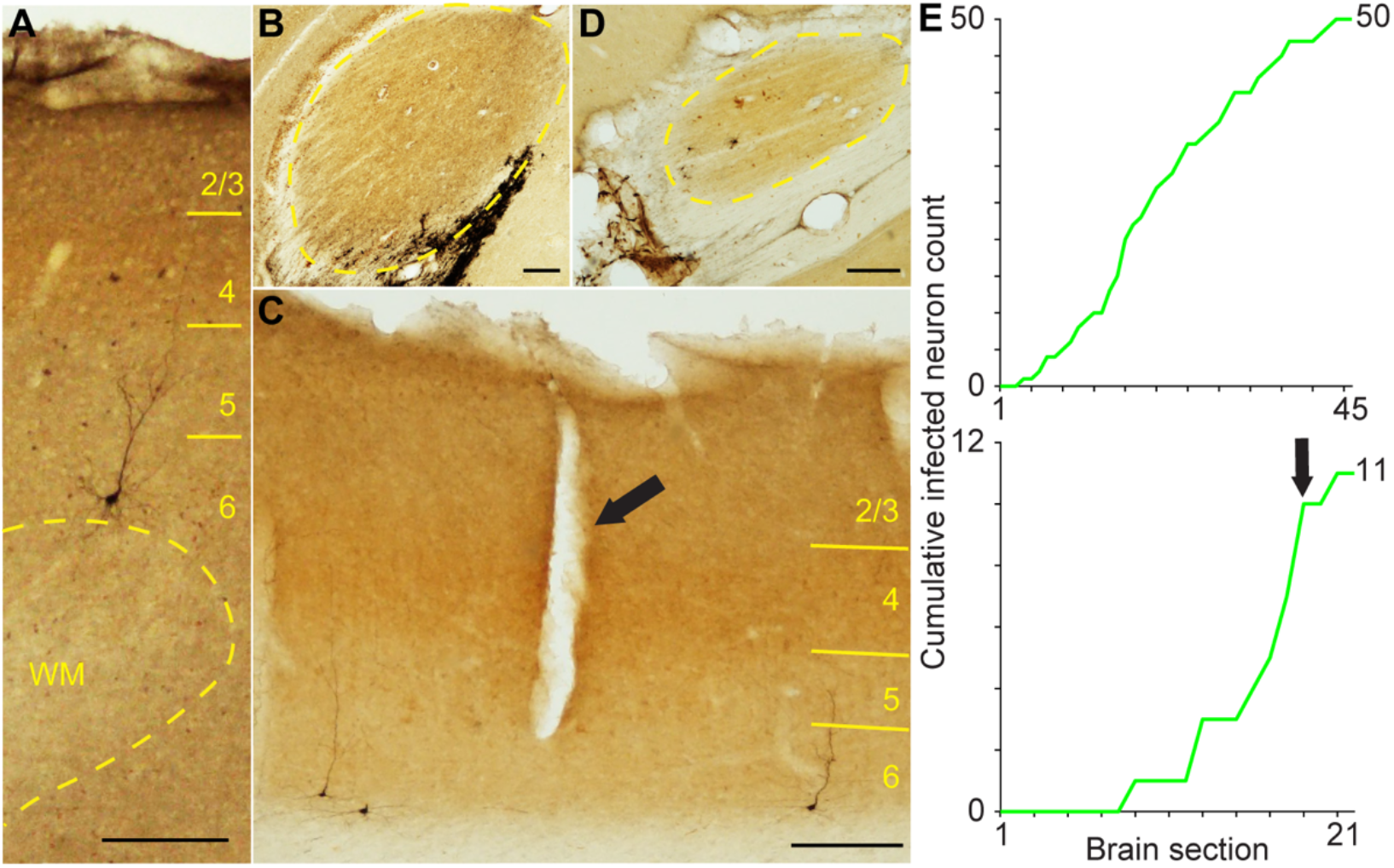
Infected corticogeniculate neurons in area 17 (V1) following virus injection into the LGN. **A** and **C**. Corticogeniculate neurons infected with Rabies-ArchT-GFP in coronal sections of area 17 of Experimental animals II and I, respectively. Sections are stained for cytochrome oxidase activity to visualize layers (labeled in yellow). Scale bars represent 200μm throughout. Arrow in **C** indicates an electrode tract. **B** and **D**. Coronal sections through LGN from the ferrets illustrated in **A** and **C**, respectively. Dark staining indicates virus infection zones and infected LGN neurons. LGN boundaries are indicated by yellow dash lines. **E**. Cumulative distributions of virus-infected neurons across coronal sections of area 17 from Experimental animals II (top) and I (bottom), respectively. Arrow indicates the section with the electrode tract shown in **C**.

### Optogenetic suppression of corticogeniculate feedback reduced LGN activity in the absence of visual stimulation

To explore the impact of corticogeniculate suppression in the absence of visual stimulation, we measured spiking activity in LGN neurons while ferrets were presented with a grey screen. Corticogeniculate feedback was optogenetically suppressed by shining a green LED (515 nm) on the surface of V1. During the first 15s following LED onset, activity among LGN neurons gradually decreased in Experimental animals (**Figure 2A, B**). This effect was quantified in two ways. First, a straight line was fit to the first 15s of the spike density function (SDF) for each neuron and the slope of this linear fit was calculated. These slopes were significantly more negative in Experimental animals compared to the Control animal (p=1.3×10^-6^, two-tailed unpaired t-test; average slope Experimental animals=-0.040±0.006, average slope Control animals=0.009±0.006; **Figure 2C**), indicating a significant reduction in LGN activity following suppression of corticogeniculate feedback. To further quantify the effect of optogenetic suppression of corticogeniculate feedback on LGN spiking activity, the area between the baseline firing rate extrapolation line and the neuronal SDF was computed over the full duration of LED presentation (denoted as the area between the curves or ABC and illustrated as the area between the grey and green curves in **Figure 2A, B**). The ABC was also significantly more negative in Experimental animals compared to the Control animal (p=0.025, two-tailed unpaired t-test; average ABC Experimental animals=-0.565±0.114, average ABC Control animals=-0.060±0.203; **Figure 2D**), further indicating that suppression of corticogeniculate feedback significantly reduced LGN spiking activity in the absence of visual stimulation. The slope was significantly more negative in Experimental animal I than Experimental animal II (p= 0.042, two-tailed unpaired t-test; average slope Experimental animal I=-0.045±0.006, average slope Experimental animal II=-0.017±0.009; **Figure 2E**). The AUC was also more negative in Experimental animal I than Experimental animal II, but this difference was not significant (p=0.394, two-tailed unpaired t-test; average slope Experimental animal I=-0.284±0.089, average slope Experimental animal II=-0.084±0.158; **Figure 2F**). Although the effect of LED illumination on LGN spontaneous firing rates was larger in Experimental animal I than that in Experimental animal II, Experimental animal I had fewer virus-infected corticogeniculate neurons per histological section of visual cortex (average number of neurons per section: Experimental animal I=0.5; Experimental animal II=1.1 neurons), so the difference in effect size was not due to infection rates. Instead, it is possible that effect sizes differed because the LED was placed closer to virus-infected corticogeniculate neurons in Experimental animal I (**Figure 1C, E**). Interestingly, firing rates among LGN neurons and multiunits were more suppressed by higher frequency (>8Hz) LED flashes compared to low frequency (<2Hz) LED flashes (p=0.0019, two-tailed paired t-test; average low frequency firing rate= 83.4±14.9Hz, average high frequency firing rate=77.3±13.9Hz). LED power at the cortical surface was higher during LED flashes greater than 8Hz compared lower frequency flashes (power >= 8Hz=0.5mW/mm^2^; power > 8Hz=0.75-1.5mW/mm^2^), thus some of this difference could be due to increased LED power at high flash frequencies. Additonally, higher frequency LED flashes induced slow oscillations in V1, described below, that could have altered LGN spontaneous firing rates. Also, frequency dependent LED-induced changes in LGN spontaneous firing rates were not due to the order of trials (lower frequency flash trials occurred before higher frequency flash trials) because an analysis limited to trials at the end of the test, after more than 30 seconds of LED-induced suppression of corticogeniculate neurons, revealed that firing rates among LGN neurons were still more suppressed by high compared to low frequency LED flash rates (p=0.002, two-tailed paired t-test).

**Figure 2:**
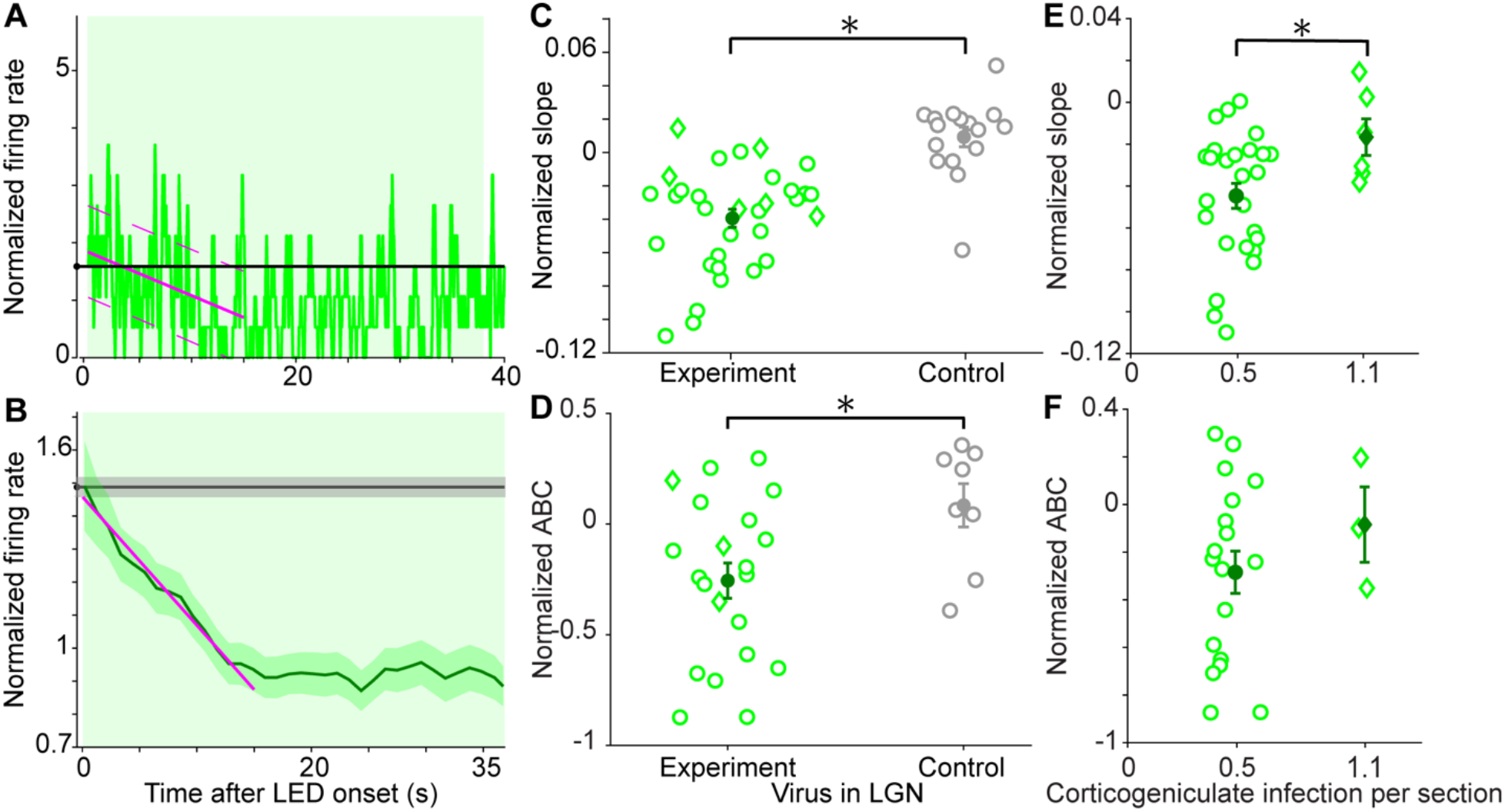
Optogenetic suppression of corticogeniculate feedback reduces firing rates among LGN neurons in the absence of visual stimulation. **A**. Spike density function (SDF) for an example LGN neuron during optogenetic suppression of corticogeniculate neurons in the absence of visual stimulation. The firing rate is normalized to the mean firing rate during the LED-ON period, indicated by green shading (0-40s). The black line represents the baseline firing rate. The solid purple line is a fit to the response during the first 15s and the dashed line is the standard error of this fit. **B**. Average and smoothed SDFs of LGN activity during optogenetic suppression of corticogeniculate feedback in the absence of visual stimulation for 19 LGN neurons recorded in 2 Experimental animals, in which ArchT was expressed in corticogeniculate neurons. Shaded areas indicate SEMs; conventions as in **A. C**. Slopes of fits to the first 15s of SDFs for 19 LGN neurons from 2 Experimental animals (32 data points, green circles for Experimental animal I and green diamonds for Experimental animal II, left) and 11 LGN neurons from 1 Control animal (16 data points, grey circles, right) in which ArchT was not expressed. The solid dots indicate average slopes for Experimental (green) and Control (grey) animals; error bars represent SEMs. Asterisk indicates significant differences in slopes across Experimental and Control animals (p=1.3 x 10^-6^, two-tailed unpaired t-test). **D**. Area between curves (ABC): baseline firing rate and firing rate during the LED-ON period for 15 LGN neurons from 2 Experimental animals (21 data points, green circles for Experimental animal I and green diamonds for Experimental animal II, left) and 6 LGN neurons from 1 Control animal (8 data points, grey circles, right). Both firing rates are normalized to the mean firing rate during the LED-ON period. Negative ABC values indicate baseline firing rate greater than LED-ON period firing rate. The solid dots indicate average ABC for Experimental (green) and Control (grey) animals; error bars represent SEMs. Asterisk indicates significant difference in ABC values across Experimental and Control animals (p=0.025, two-tailed unpaired t-test). **E.** Slopes from Experimental animals in **C** were plotted against the average number of infected corticogeniculate neurons per histology section of visual cortex, separately for 13 LGN neurons from Experimental animals I (26 data points, green circles, left) and 6 LGN neuronos from Experimental animals II (6 data points, green diamonds, right). Asterisk indicates significant differences in slopes across Experimental animals (p= 0.042, two-tailed unpaired t-test). Conventions as in **C. F.** ABCs from Experimental animals in **D** were plotted against the average number of infected corticogeniculate neurons per histology section of visual cortex, separately for 12 LGN neurons from Experimental animals I (18 data points, green circles, left) and 3 LGN neurons from Experimental animals II (3 data points, green diamonds, right). Conventions as in **D**.

### Optogenetic suppression of corticogeniculate feedback did not affect LGN visual responses

We next examined whether the visual responses of LGN neurons were affected by optogenetic suppression of corticogeniculate feedback. Visual responses were recorded during presentations of drifting gratings varying in contrast, spatial frequency, temporal frequency, or size, with optogenetic suppression of corticogeniculate feedback induced by LED illumination of V1 on half of the trials. LGN neurons were classified as X or Y if they demonstrated higher/lower c50 values, lower/higher preferred temporal frequencies, and little/more surround suppression, respectively (**Table 1**), consistent with known physiological differences among carnivore X and Y LGN neurons (Derrington & Fuchs 1979). In contrast to the impact of optogenetic suppression of corticogeniculate feedback on LGN activity in the absence of visual stimulation, LGN responses to visual stimuli were largely unaffected by optogenetic suppression of corticogeniculate feedback. **Figure 3A** illustrates the average PSTH of LGN neurons that were tuned for changes in stimulus contrast, recorded in Experimental animals. The average PSTH illustrates two repeats of stimuli increasing in contrast during LED suppression of corticogeniculate feedback (**Figure 3A**, green bars indicate periods of LED illumination). We again quantified the slope of the linear fit to the first 15s of each neuronal PSTH (as in **Figure 2A**). As illustrated in the average PSTH of LGN neuronal responses to gratings increasing in contrast, the slope of the fit to the first 15s of neuronal responses was flat (**Figure 3A**), suggesting no reduction in LGN responses to visual stimulation during optogenetic suppression of corticogeniculate feedback. In order to quantify the effect of optogenetic suppression of corticogeniculate feedback on LGN responses to visual stimulation, we compared slopes of LGN neuronal PSTHs for each visual stimulus type on trials without and with LED illumination of V1. This comparison was made only for LGN neurons recorded in Experimental animals. We observed no significant difference in the slope of LGN neuronal responses to any type of visual stimulation across LED conditions (p>0.38 for all, two-tailed paired t-test; **Table 2**) and slopes were mainly clustered around zero for both LED conditions indicating flat linear fits (**Figure 3C**). To further examine the effects of optogenetic suppression of corticogeniculate feedback on LGN visual responses, we compared the magnitudes of LGN neuronal responses to visual stimuli as well as tuning curve metrics across LED conditions. **Figure 3B** illustrates spatial frequency and contrast tuning curves for a representative LGN neuron recorded in an Experimental animal across both LED conditions. There is little change in the magnitude of this example neuron’s response across LED conditions, nor does the neuron’s preferred tuning change. To quantify neuronal response magnitude, we computed the area under the curve (AUC) for each tuning curve for each LGN neuron recorded in Experimental animals. As was the case for the representative example, LGN response magnitude quantified as the AUC did not differ across LED conditions for any visual stimulus type displayed (p>0.17 for all, twotailed paired t-test; **Figure 3D, Table 2**). Additionally, we observed no changes in LGN tuning metrics across LED conditions (**Table 1**). The only exception to the overall lack of an effect of optogenetic suppression of corticogeniculate feedback during visual stimulus presentation was a small, but significant, reduction in the maximum evoked firing rate of LGN neurons during spatial frequency tuning tests with LED suppression of feedback (p=0.012, paired t-test with Bonferroni corrected alpha=0.0125; maximum firing rate no LED=15.2±2Hz, maximum firing rate with LED=11.6±1.3Hz). No differences in maximum evoked firing rates were observed for other tuning tests (p>0.06, paired t-test with Bonferroni corrected alpha=0.0125). Together, these results suggest that while optogenetic suppression of corticogeniculate feedback reduces LGN firing rates, visual stimulation appears to override this suppression.

**Figure 3:**
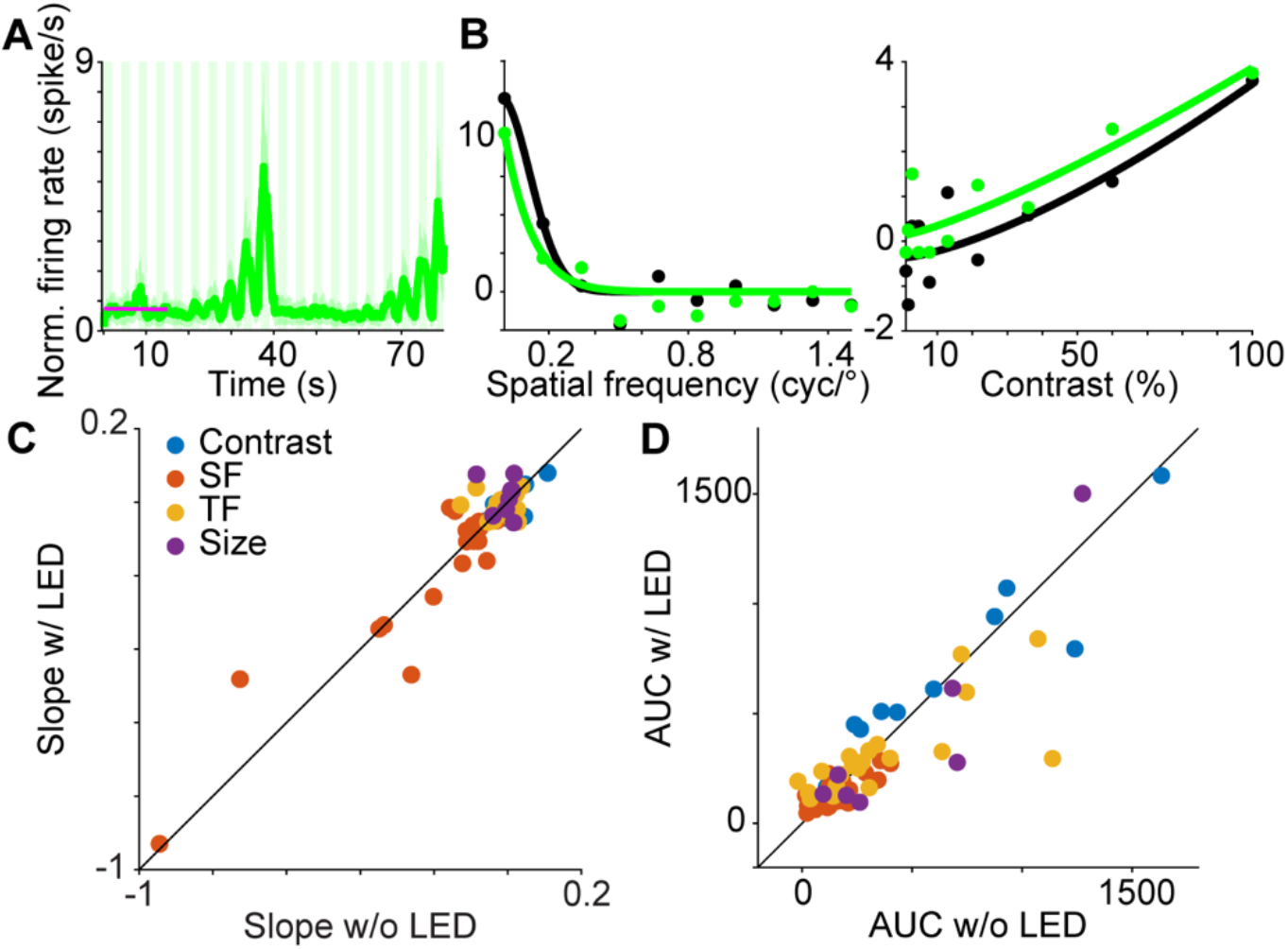
Optogenetic suppression of corticogeniculate feedback does not alter visual response properties of LGN neurons. **A**. Average smoothed peri-stimulus time histogram (PSTH) of 11 LGN neurons from 2 Experimental animals in response to two repeats of gratings increasing in contrast. The solid purple line is a fit to the response during the first 15s. **B**. Spatial frequency (left) and contrast (right) tuning curves for an example LGN neuron recorded in an Experimental animal. Black and green curves are fits to data (dots) on trials without and with optogenetic suppression of corticogeniculate feedback, respectively. **C**. Slopes of fits to the first 15s per PSTH in response to gratings varying in contrast (blue), spatial frequency (orange), temporal frequency (yellow) and size (purple) on trials without and with LED-induced suppression of corticogeniculate neurons. See **Table 2** for statistics. **D**. Area under curves (AUCs) for baseline-subtracted tuning curves, on trials without and with LED-induced suppression of corticogeniculate neurons. Tuning data color-coded as in **C**. See **Table 2** for statistics.

Activation of corticogeniculate feedback reduces visual response latencies and increases visual response precision among LGN neurons (Hasse & Briggs 2017). We therefore asked whether suppression of corticogeniculate feedback might also affect the timing of LGN visual responses. We quantified LGN visual response latency in two ways. First, we measured visual response latency as the time between the preferred stimulus and the spike recorded during presentation of m-sequence white noise stimuli. As illustrated by a representative LGN neuron (**Figure 4A**), and the sample (**Figure 4B**), most LGN neurons (6 of 10) showed an increased response latency during LED-induced suppression of corticogeniculate feedback, although this effect was not significant (p=0.164, two-tailed paired t-test; average latency no LED=10.8±3.90, average latency with LED=15.6±5.04). As a secondary confirmation of the effect of optogenetic suppression of corticogeniculate feedback on LGN response latency, we also measured LGN latencies and spike timing precision in response to a flashing spot stimulus. We observed no change in response latency across LED conditions for four LGN neurons with clear responses to the flashing stimulus (p=0.789, two-tailed paired t-test; average latency no LED=24.8±1.49 msec, average latency with LED=25.0±1.73 msec; **Figure 4C**). Likewise, spike timing precision, quantified as the standard deviation (STD) of the time of the first spike in response to a flash, was not significantly different across LED conditions (p=0.847, two-tailed paired t-test; average STD first spike time no LED=8.47±1.48 msec, average STD first spike time with LED=8.04±1.64 msec; **Figure 4D**). Thus, suppression of corticogeniculate feedback did not significantly influence spike timing in the LGN, although there was a trend toward an increase in LGN response latencies during presentation of m-sequence stimuli paired with optogenetic suppression of corticogeniculate feedback.

**Figure 4:**
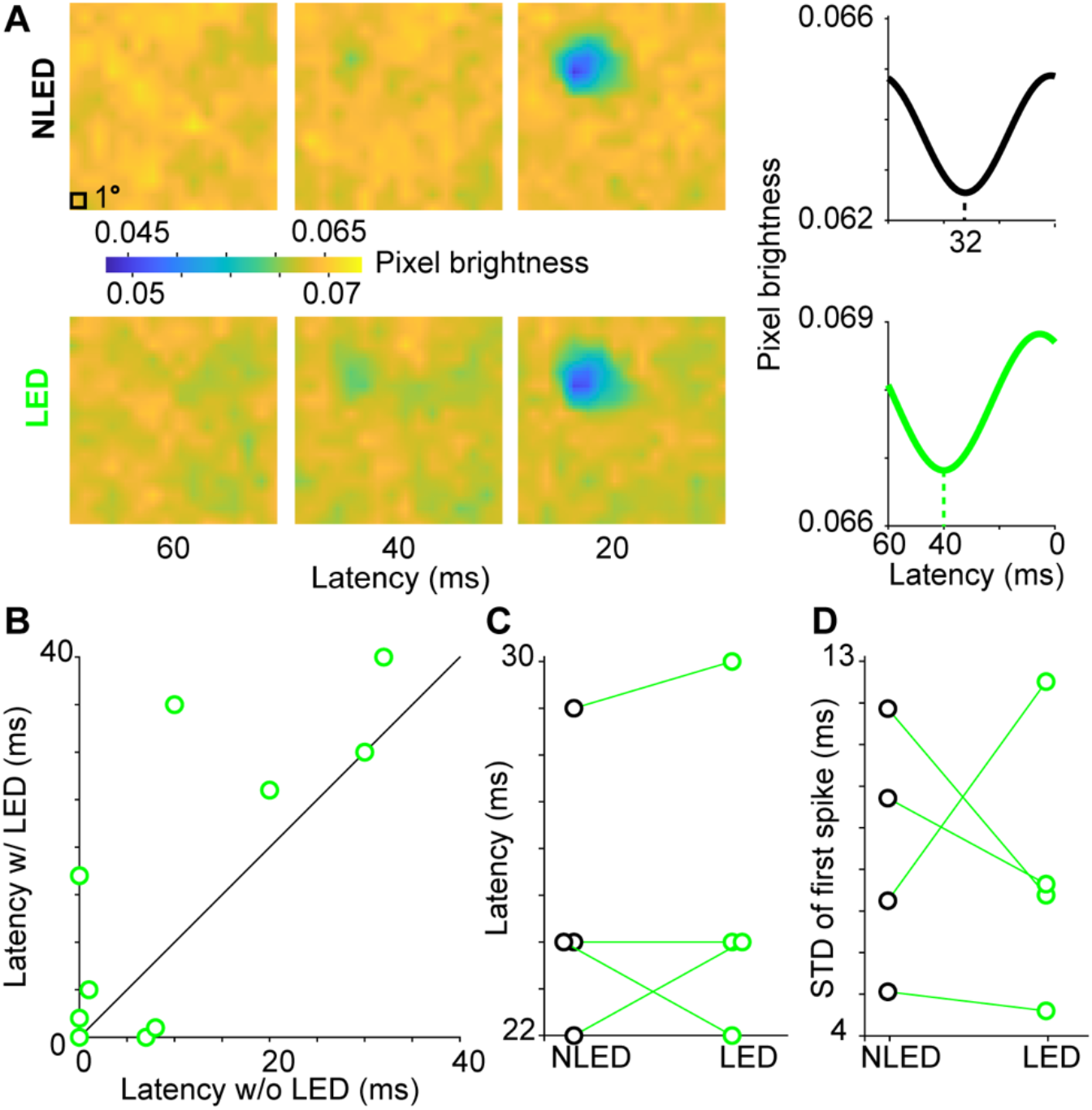
Optogenetic suppression of corticogeniculate feedback does not alter visual response latencies or spike timing precision among LGN neurons. **A**. Left: spatiotemporal receptive fields of a representative LGN OFF neuron recorded in an Experimental animal without (top row) and with (bottom row) LED-induced suppression of corticogeniculate neurons. Color bar represents pixel brightness; black box illustrates 1 degree. Right: pixel brightness curves measured from the spatiotemporal receptive fields at left. Black and green curves are without and with LED-induced suppression of corticogeniculate neurons, respectively. Dashed black and green lines illustrate calculation of response latency. **B**. Response latencies of 10 LGN neurons from 2 Experimental animals measured from spatiotemporal pixel brightness curves on trials without and with LED-induced suppression of corticogeniculate neurons. **C**. Response latencies of 4 LGN neurons from 2 Experimental animals in response to a flashed stimulus on trials without (black) and with (green) LED-induced suppression of corticogeniculate neurons. **D**. Standard deviations of the first spike time for 4 LGN neurons from 2 Experimental animals in response to a flashed stimulus. Conventions as in **C**.

### LED illumination shifts oscillations in V1 spiking patterns

Results thus far suggested that optogenetic suppression of corticogeniculate feedback reduces the firing rates of LGN neurons, but these effects are overridden when strong visual stimulus drive arrives through the feedforward retinogeniculate pathways. We next explored changes in V1 neuronal activity during optogenetic suppression of corticogeniculate neurons that might support the observed effects in the LGN. **Figure 5A** illustrates 30s of V1 neuronal responses to LED flashes at varying frequencies in the absence of visual stimulation among four well isolated neurons simultaneously recorded in the deep layers of V1 in Experimental animal II. There were fewer spikes during low-frequency LED flash trials (e.g., first row in **Figure 5A**) compared to higher-frequency LED flash trials (e.g., second row in **Figure 5A**). Firing rates among these four V1 neurons were lower during low-frequency (0.5-1.5Hz) compared to high-frequency (8.5-16.5Hz) LED flashes (mean spike rate low=2.51±0.791 spikes/s; mean spike rate high=3.52±1.54 spikes/s; **Figure 5C**), although this difference did not reach statistical significance due to the limited sample size (p=0.0783, two-tailed paired t-test). In contrast, a larger sample of V1 multiunits recorded from a wider range of V1 deep layer contacts did not show a significant difference in firing rate during low-versus high-frequency LED flashes (p=0.831, two-tailed paired t-test; mean spike rate low=14.2±1.66 spikes/s; mean spike rate high=14.3±1.63 spikes/s; **Figure 5C**). Based on the patterns observed among the V1 neurons, we hypothesized that LED illumination of V1 shifted spiking patterns among V1 neurons depending on the frequency of LED flashes. To test this hypothesis, we computed power spectral density curves for all V1 single- and multiunits during low- and high-frequency LED flashes in the absence of visual stimulation. While low-frequency LED flashes enabled V1 units to display a variety of oscillatory patterns (**Figure 5B**, left), high-frequency LED flashes shifted all V1 units toward similar low-frequency oscillations (**Figure 5B**, right). Accordingly, peak oscillation frequencies per V1 unit were distributed in response to low-frequency LED flashes, but were tightly clustered at lower frequencies in response to high-frequency LED flashes (**Figure 5D**). Indeed, peak oscillation frequencies among V1 units during high-frequency LED flashes were significantly lower than those observed during low-frequency flashes (p=0.0251, two-tailed paired t-test; mean peak frequency low=0.296±0.028Hz, mean peak frequency high=0.209±0.012Hz). Additionally, there was significantly less variation in the peak oscillation frequency among V1 units in response to high-versus low-frequency LED flashes (p=0.0018, F test). These findings suggest that optogenetic suppression of corticogeniculate neurons using medium-to high-frequency LED flashes, as utilized for most tests, shifted V1 neuronal spiking activity into a 0.2Hz oscillation that promoted net suppression of spiking activity in the LGN. Indeed net suppression of LGN activity was greater during high-compared to low-frequency optogenetic suppression of corticogeniculate feedback.

**Figure 5:**
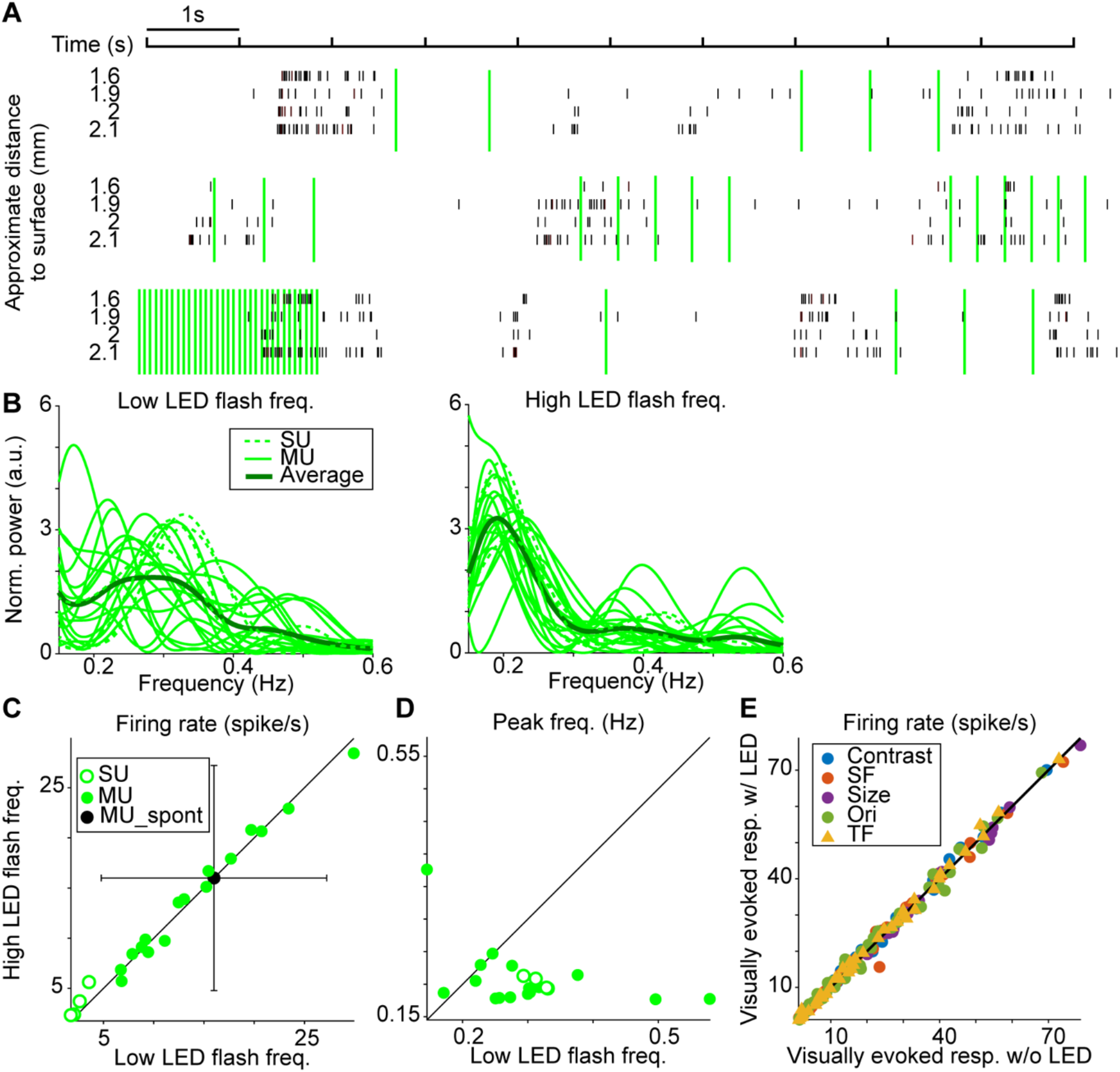
Effects of optogenetic suppression on deep layer V1 neurons. **A**. Spike raster of four V1 deep layer neurons recorded simultaneously in an Experimental animal in response to LED flashes (green lines) at varying frequencies. **B**. Normalized power spectra of 4 neurons/single units (dashed green lines; SU) and 16 multiunits (solid green lines; MU) recorded in V1 of one Experimental animal during low-frequency (0.5-1.5Hz, left) and high-frequency (8.5-16.5Hz, right) LED flashes. Power spectra were each normalized to the averaged power between 0.15 and 0.6Hz. Thicker dark green lines indicate average power spectra for V1 units. **C**. Firing rates of V1 SUs (open green circles) and MUs (filled green circles) during low- and high-frequency LED flashes. Filled black dot indicates the spontaneous firing rate of MUs, measured during the first 250 msec of recordings prior to LED flashes; error bars show SD. **D**. Frequencies corresponding to peak power among V1 SUs and MUs during low- and high-frequency LED flashes; conventions as in **C. E**. Average visually evoked firing rates among 40 V1 MUs from two Experimental animals during measurements of contrast, spatial frequency, temporal frequency, and size tuning on trials without and with LED suppression of corticogeniculate neurons. Tuning data color-coded as in **Figure 3C**. See **Table 3** for statistics.

Studies of corticothalamic neurons in mice suggest that corticothalamic neurons are locally connected to inhibitory neurons in V1 that suppress the activity of other V1 neurons (Bortone et al 2014, Olsen et al 2012). We hypothesized that higher frequency LED-induced suppression of corticogeniculate neurons may lead to dis-inhibition of neighboring V1 neurons, in addition to shifting V1 spiking into a 0.2Hz oscillation. To test this notion, we examined the responses of V1 multiunits to drifting gratings varying in temporal frequency. Importantly, because the LED flash frequency was coupled to the temporal frequency of the drifting grating, many trials of the temporal frequency tuning test included high-frequency LED flashes. In support of the dis-inhibition hypothesis, we observed a small but significant increase in the firing rate of V1 multiunits in response to gratings varying in temporal frequency on trials with compared to without LED illumination of V1 (p=0.0026, two-tailed paired t-test; **Figure 5E**, **Table 3**). Interestingly, there were no significant differences in firing rate among V1 multiunits across LED conditions for other tuning tests, where the LED flash frequency was fixed at 4Hz (p>0.02 for all, Bonferroni corrected alpha=0.01; **Figure 5E**, **Table 3**).

**Table 3:**
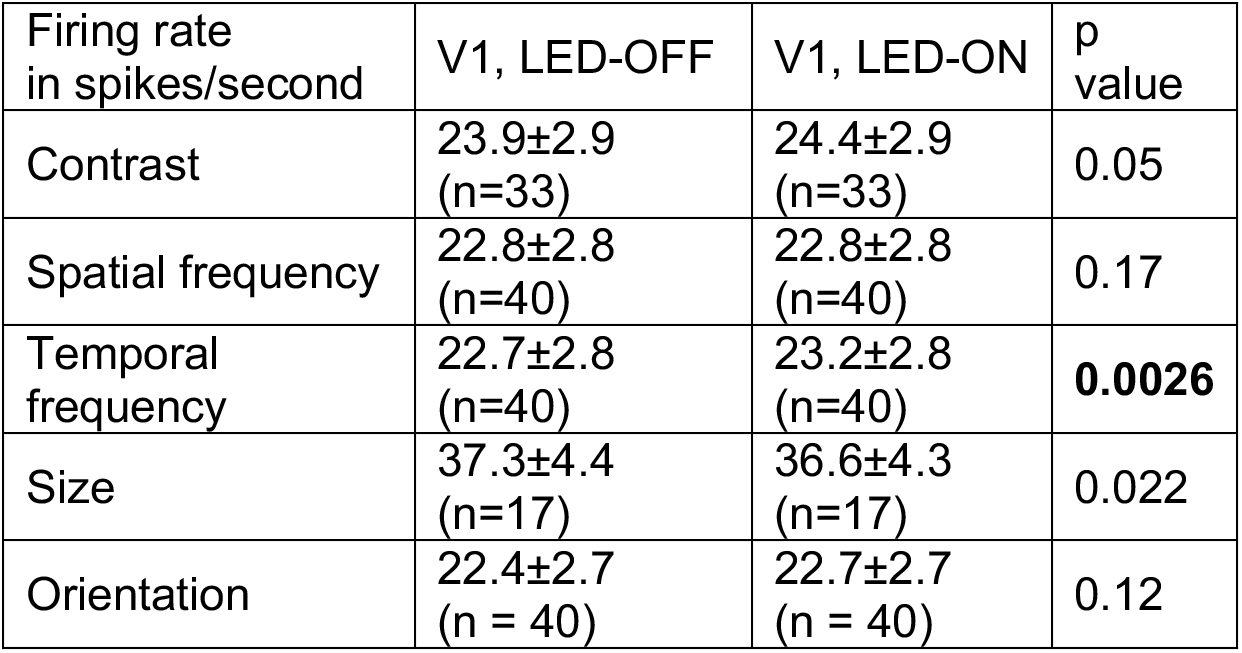
Effects of optogenetic suppression of corticogeniculate neurons on V1 multiunit firing rate during visual stimulation. P values were calculated from paired t-test across LED conditions and corrected for multiple comparisons using Bonferroni correction (alpha=0.01).

## Discussion

Although many studies have investigated the role of corticogeniculate feedback in visual perception, measured effects are often subtle and findings have not necessarily been consistent across studies, leading to a lack of overall consensus on the function of corticogeniculate feedback. Subtle and/or inconsistent effects across studies could be due to a number of factors including the fact that corticogeniculate circuits are heterogeneous (Briggs 2020) and their influence may vary depending on their mode of activation (Kirchgessner et al 2020) and may be overridden by strong feedforward visual stimulus drive (Guillery & Sherman 2002). One major concern is that corticothalamic circuits are suppressed during slow cortical oscillations in anesthesia and sleep (Steriade 2003), so suppression of corticogeniculate feedback in anesthetized preparations may run into a floor effect. Another major concern is that non-selective and irreversible manipulations that impact V1 neurons generally, in addition to corticogeniculate neurons, may confound interpretation of results and make it difficult to quantify whether a floor effect is present with anesthesia. Accordingly, it is critical to address whether methodological differences or any of these confounding factors contribute to inconsistent findings on corticogeniculate influence over LGN activity. We examined whether selective and reversible optogenetic suppression of corticogeniculate feedback influenced LGN neuronal activity with and without visual stimulation and under different modes of optogenetic suppression, all in anesthetized ferrets. These experiments allowed us to explicitly test whether anesthesia produced a floor effect, whether LGN and V1 neuronal responses varied under different modes of corticogeniculate suppression, and whether visual stimulus drive or other confounding factors contributed to results. Somewhat surprisingly, we observed a reduction in LGN spiking activity when corticogeniculate feedback was optogenetically suppressed (**Figure 2**), demonstrating that anesthesia does not induce a complete floor effect. Additionally, higher versus lower frequency optogenetic suppression of corticogeniculate neurons differentially affected the activity of V1 and LGN neurons: high frequency optogenetic suppression of corticogeniculate neurons shifted V1 neurons into a 0.2Hz oscillation (**Figure 5**), relaxed local inhibition within V1, and generated more net suppression of LGN activity. Frequency-dependent optogenetic effects could have been due in part to higher LED power at higher frequency illumination, but suppression of LGN spontaneous firing rates was present for all LED flash frequencies and visual stimulation overroad optogenetic suppression regardless of LED frequency. Most revealing was our discovery that the net inhibitory influence of corticogeniculate suppression on LGN activity was completely overridden by feedforward visual stimulus drive. In other words, although spiking activity among LGN neurons was consistently reduced during optogenetic suppression of corticogeniculate feedback in the absence of visual stimulation (**Figure 2**), visually driven responses and tuning metrics for LGN neurons were mostly unchanged when corticogeniculate feedback was suppressed (**Figures 3, 4, Tables 1, 2**). Together these findings show that selective and reversible suppression of corticogeniculate neurons produces clear, measurable, and consistent effects in both the LGN and locally in V1, even in anesthetized animals. However, corticogeniculate influence is subtle and easily overridden by feedforward drive generated by the display of visual stimuli that strongly modulate LGN neuronal activity in highly visual animals.

Importantly, our findings provide some explanations for inconsistencies in previous studies. For example, a number of studies have reported different effects of non-selective suppression of corticogeniculate feedback on response gain, adaptation, and temporal modulation for LGN neurons in distinct parallel streams (Gulyas et al 1990, Li et al 2011, Przybyszewski et al 2000). In all of these cases, effects of feedback suppression were observed when LGN responses to non-preferred stimuli were measured (e.g. magnocellular and Y cell responses to high-contrast stimuli). Our results suggest suppression of corticogeniculate feedback may only be visible when feedforward visual stimulus drive is reduced, as is the case when LGN neurons are poorly driven by non-preferred stimuli. Along similar lines, Wang et al (2018) observed diverse effects of non-selective activation of corticogeniculate feedback on LGN responses to flashing probe stimuli that are very good at driving retinal ganglion cells, but do not generate strong responses among V1 neurons, including corticogeniculate neurons. Accordingly, subtle influences of corticogeniculate feedback were likely overridden by strong feedforward visual stimulus drive in this case. Similarly diverse effects of selective and non-selective corticogeniculate suppression have also been observed in anesthetized mice and cats, respectively (Denman & Contreras 2015, Geisert et al 1981). In both of these cases, spontaneous firing rates among recorded LGN neurons were low, making it difficult to assess effects of corticogeniculate suppression on LGN activity in the absence of visual stimuli. And in both cases, a variety of visual stimuli produced a range of responses among LGN neurons, some demonstrated changes during corticogeniculate suppression and some not. It is unclear whether a deeper examination of LGN responses to preferred and non-preferred stimuli would reveal more consistent effects of corticogeniculate suppression in these studies.

Recent evidence suggests that sensory thalamic responses vary depending on the firing mode of corticothalamic neurons, introducing another potential source of variation across studies of corticogeniculate feedback. Optogenetic stimulation of corticothalamic neurons at 10Hz in awake mice and in brain slices result in net facilitation of LGN and ventral posteromedial thalamus (VPM) activity, while 0.1Hz stimulaton in brain slices and continuous stimulation in awake mice result in net suppression of sensory thalamic neurons (Crandall et al 2015, Denman & Contreras 2015, Kirchgessner et al 2020). In this study, we suppressed, rather than activated, corticogeniculate neurons, but we observed some interesting changes in LGN and V1 activity that complement observations from prior work. Specifically, when we optogenetically suppressed corticogeniculate neurons at frequencies near 10Hz (~8-17Hz), V1 neurons synchronized their activity into a 0.2Hz oscillation (**Figure 5B, D**) and LGN spiking activity was more suppressed. In contrast, lower frequency optogenetic suppression of corticogeniculate neurons silenced some putative corticogeniculate neurons (**Figure 5A**), but did not systematically shift V1 population activitiy (**Figure 5B, C**), and resulted in slightly less suppression of LGN activity. These findings suggest that both optogenetic activation and suppression of corticogeniculate neurons at different frequencies generate different effects within V1 and the LGN. Furthermore, optogenetic activation/suppression of corticogeniculate neurons around 10Hz generates complementary net facilitation/suppression in the LGN, respectively. Together these findings also suggest that the precise temporal control of corticogeniculate neurons offered by optogenetics may be important when studying corticogeniculate contributions to LGN activity.

Hasse & Briggs (2017) discovered that optogenetic activation of corticogeniculate feedback in anesthetized ferrets improves the temporal precision of LGN neuronal responses to visual stimuli, including reducing their average response latencies to both grating and m-sequence stimuli. In a follow-up study, Murphy et al (2020) demonstrated that the improvement in temporal resolution was due to a reduction in LGN neuronal response variability when corticogeniculate feedback was optogenetically activated. We hypothesized that optogenetic suppression of corticogeniculate feedback might increase LGN response variability and increase average response latencies. Although we observed a trend whereby response latencies during m-sequence presentation increased for the majority of LGN neurons (**Figure 4A, B**), our sample was not sufficiently large to reach statistical significance. Interestingly, we did not observe a change in latency or spike timing precision for LGN neuronal responses to gratings when corticogeniculate feedback was optogenetically suppressed (**Figure 4C, D**), nor did we observe changes in LGN response variability (data not shown). However, given our main finding that weak effects of corticogeniculate suppression are overridden by feedforward visual stimulus drive, these observations make more sense. M-sequence stimuli drive LGN neurons more sparsely than gratings. Thus suppressive effects were more apparent during m-sequence presentation compared to grating presentation, when LGN responses were dictated more strongly by feedforward visual stimulus drive.

Selective and reversible manipulation of corticogeniculate neurons through optogenetics can reveal not only corticogeniculate influence on LGN activity, but also corticogeniculate influence locally among other V1 neurons. Non-selective and irreversible manipulations of V1 do not permit this type of examination. In mice, continuous activation of corticothalamic neurons suppresses neuronal activity throughout V1 via activation of local inhibitory neurons (Bortone et al 2014, Olsen et al 2012). We therefore asked whether optogenetic suppression of corticogeniculate neurons might relax local inhibition within V1, generating an increase in response gain among V1 neurons. Interestingly, we observed a significant increase in V1 neuronal response gain to gratings varying in temporal frequency when corticogeniculate neurons were optogenetically suppressed (**Figure 5E, Table 3**). Importantly, the LED was synchronized to the visual stimulus grating cycle, meaning that many trials of the temporal frequency tuning test involved high-frequency optogenetic suppression of corticogeniculate neurons. These findings support the notion that corticogeniculate neurons are coupled to local inhibitory neurons that inhibit other V1 neurons and high-frequency activation/suppression of corticogeniculate neurons adjusts this local gain control mechanism. It is also noteworthy that we observed a synchronization of V1 activity into a 0.2 Hz oscillation when corticogeniculate neurons were optogenetically suppressed using 8.5-16.5Hz LED flashes. Corticothalamic neurons generate slow oscillations (<1Hz) intracortically during anesthesia and non-REM sleep (Steriade 2003), and high-frequency optogenetic suppression of corticogeniculate neurons in this study entrained this oscillation more. Here we document that in addition to V1 slow oscillations, high-frequency optogenetic suppression of corticogeniculate neurons also induces a prolonged suppression of LGN spiking activity that lasts many seconds (**Figure 2**). Together these patterns of activity could be important for coordinating corticothalamic interactions in sleep.

In summary, we find that selective and reversible suppression of corticogeniculate feedback causes clear and consistent effects on LGN and V1 neuronal activity, even under anesthesia, dispelling the notion that anesthesia produces a floor effect. More importantly, we demonstrate that weak influences of corticogeniculate suppression are easily overcome by strong feedforward visual stimulus drive. And this weak influence of feedback relative to strong feedforward drive is the most likely source of inconsistencies in the literature. Additionally, as more information becomes available using selective and reversible manipulation methods, it is clear that corticogeniculate function must be studied using methods that offer precise temporal control, as the influence of corticogeniculate feedback on LGN and V1 neuronal activity depends on the activity regime that corticogeniculate neurons are in at any given moment. These methods also enable examination of corticogeniculate influence over local activity within V1, an increasingly important avenue for further exploration.

## Author Contributions

S.Z., J.M.H, and F.B. designed the research. J.M.H. and F.B. collected the data. S.Z. analyzed the data. S.Z. and F.B. wrote the manuscript.

## Acknowledgements

We thank Elise Bragg for expert technical assistance and Drs. Karen Moodie and Kirk Maurer for veterinary assistance. This work was funded by National Institutes of Health (National Eye Institute: EY018683 and EY025219 to F.B.), National Science Foundation (EPSCoR 1632738), the Whitehall Foundation, the Hitchcock Foundation, the Del Monte Institute for Neuroscience at the University of Rochester, and a University of Rochester University Research Award. J.M.H. was supported by a Graduate Fellowship from the Albert J. Ryan Foundation.

